# Interactions between thermal and nutritional stress reveal cryptic adaptive potential

**DOI:** 10.64898/2026.10.18.759992

**Authors:** Greg M. Walter, Avishikta Chakraborty, Luke Yates, Fiona Cockerell, Craig R. White, Carla M. Sgrò

**Affiliations:** School of Biological Sciences, Monash University, Melbourne 3800, Australia; School of Natural Sciences, University of Tasmania, Hobart 7005, Australia; Department of Genetics, Evolution & Environment, University College London, London WC1E 6BT, UK

**Keywords:** adaptive potential, diet stress, cryptic genetic variation, *Drosophila*, fitness, fundamental theorem of natural selection, genotype-by-environment interactions, G×E×E, evolutionary rescue, productivity

## Abstract

Predicting whether populations can adapt fast enough to persist under global change requires understanding how adaptive potential changes with the environment. Theory predicts that cryptic genetic variation hidden under benign conditions may be released under stress, but empirical tests are rare and typically manipulate single stressors in isolation, which is not realistic under global change. Using a quantitative genetic breeding design with *Drosophila melanogaster*, we quantified additive genetic variance in fitness (female productivity) across combinations of temperature and nutritional stress. We found that nutritional stress increased genetic variance at cold and hot margins. Temperature and diet also interacted to shape which genotypes performed best: nutritional stress weakened genetic correlations across temperatures at the hot margin, whereas genetic correlations across diets strengthened at the cold margin. Adaptive potential is therefore environment-dependent and multidimensional, meaning single-stressor experiments are likely to systematically misrepresent adaptive capacity under realistic global change scenarios.

## Introduction

Global change is increasingly exposing populations to combinations of stressors (e.g., warming alongside resource limitation), which threaten their persistence (Diffenbaugh & Field 2013; Cassia *et al*. 2018). While migration and phenotypic plasticity can help populations track environmental variation, these responses often fail in conditions outside their historical experience, leaving adaptation as the primary route to persistence (Urban *et al*. 2024; Chevin & Bridle 2025). The capacity to adapt depends on the amount of additive genetic variance in fitness, the heritable variation in reproductive success that natural selection can act upon (Fisher 1930; Orr 2007, 2009). However, genetic variance in fitness can change with the environment, making it difficult to understand how global change will influence the adaptive potential of populations (Hoffmann & Merilä 1999; Charmantier & Garant 2005). In particular, we have a poor understanding of how combinations of stressors shape adaptive potential, limiting our ability to predict whether populations can adapt to realistic, multidimensional global change (Hoffmann & Sgrò 2011; Urban *et al*. 2024).

Within populations, genotypes often vary in their response to environmental change, creating genotype-by-environment interactions (G×E) that determine how adaptive potential changes with the environment (Sheth *et al*. 2018; Kulbaba *et al*. 2019; Walter *et al*. 2026b). Two forms of G×E influence adaptive potential. Firstly, G×E can change the amount of genetic variance in fitness across environments, which determines the capacity for adaptation to any given environment. Secondly, G×E is captured by genetic correlations across environments, which reflect the extent to which the same genotypes are favoured across environments, or whether different genotypes perform relatively better as conditions change (Falconer 1952; Via & Lande 1985; Sgrò & Hoffmann 2004; Saltz *et al*. 2018; Lasky *et al*. 2023). Despite the importance of G×E in shaping adaptive potential, we lack empirical tests of how additive genetic variance and genetic correlations in fitness respond to combinations of environmental stressors. This is largely due to the considerable challenge of estimating fitness for a large number of genotypes in multiple environments.

Most estimates of genetic variance in fitness come from benign laboratory conditions, or from populations in their native environments that are presumed to be near their evolutionary equilibrium (Houle 1992; Hendry *et al*. 2018; Bonnet *et al*. 2022). Yet adaptive potential matters most when populations face stressful, novel, or rapidly changing conditions that cause fitness declines and threaten persistence (Lande 1988; Gomulkiewicz & Holt 1995; Bell & Gonzalez 2009). Cryptic genetic variation that remains hidden in benign environments is expected to be expressed under stressful conditions (Le Rouzic & Carlborg 2008; McGuigan & Sgrò 2009; Paaby & Rockman 2014), which should increase genetic variance in fitness and therefore adaptive potential for populations facing decline (Gomulkiewicz & Shaw 2013; Shaw & Shaw 2014). While studies have shown that novel or stressful environments can increase adaptive potential (Parsons 1987; Sgrò & Hoffmann 1998; Torres-Martínez *et al*. 2019; Walter *et al*. 2023; Walter *et al*. 2026b), they focus on single stressors in isolation, or involve field environments in which multiple stressors are confounded.

The type and intensity of stressors are increasing under global change, which means that populations rarely face single environmental challenges in isolation (Diffenbaugh & Field 2013; Cassia *et al*. 2018; Cuenca-Cambronero *et al*. 2021). In particular, thermal stress is increasingly combined with altered precipitation patterns that influence resource availability and create novel combinations of abiotic and biotic conditions (Crain *et al*. 2008; Todgham & Stillman 2013; Cross *et al*. 2015; Rosenblatt & Schmitz 2016; Hardison & Eliason 2024). Evidence suggests that genotypes can vary in response to combinations of stressors, creating G×E×E (Des Marais *et al*. 2017; Kutz *et al*. 2019; Hudak & Dybdahl 2023). If combinations of thermal and nutritional stress expose genetic variance that was not observed under either stressor alone, then estimates of adaptive potential from single-stressor studies are likely to be misleading (Kutz *et al*. 2019; Choy *et al*. 2024), and current approaches for predicting evolutionary responses may systematically misrepresent the adaptive capacity of populations.

Environmental stress that creates negative population growth will reduce mean fitness 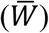 of the population. If the population is to recover mean fitness, it will need to adapt, which requires additive genetic variance in relative fitness, defined as the ratio of genetic variance in absolute fitness to mean fitness squared, 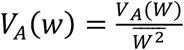 (Fisher 1930; Orr 2007, 2009; Kulbaba *et al*. 2019). Here, we test how combinations of environmental stressors shape adaptive potential by estimating additive genetic variance in relative fitness (*V*_*A*_(*w*)) and additive genetic correlations across factorial combinations of temperature and nutritional stress in a large, outbred population of *Drosophila melanogaster*. By quantifying how

*V*_*A*_(*w*) and genetic correlations change from benign to combinations of stressors, we test whether adaptive potential under combined stress can be predicted from single-stressor treatments. Our approach provides a direct test of whether multidimensional environmental change alters the magnitude and genetic basis of adaptive potential, with implications for forecasting evolutionary responses under global change.

## Material and methods

We estimated *V*_*A*_(*w*) in six treatments that combined three temperatures (14, 25 and 28°C) with two diets (standard and a low-calorie 25% dilution) (**Fig. 1**). We chose these temperatures as they span the natural thermal range of *D. melanogaster* in the wild, with 25°C representing control conditions that populations were established in, and 28°C the hot margin predicted under climate change or heat wave conditions (Choy *et al*. 2024). Nutritional stress (low-calorie diet) represents resource limitation arising from accelerated fruit decomposition, phenological mismatches, and altered nutrient content expected under climate change (Cross *et al*. 2015; Rosenblatt & Schmitz 2016). To estimate *V*_*A*_(*ω*), we used a half-sibling breeding design by mating 150 sires each to five dams, and then reared offspring from each of the resulting 750 full-sibling families in each of the six treatments (**Fig. 1**). We then measured productivity in female offspring, which captures fitness as the combination of fecundity and larval viability (Mackay 1985; Nguyen & Moehring 2015).

**Fig. 1.**
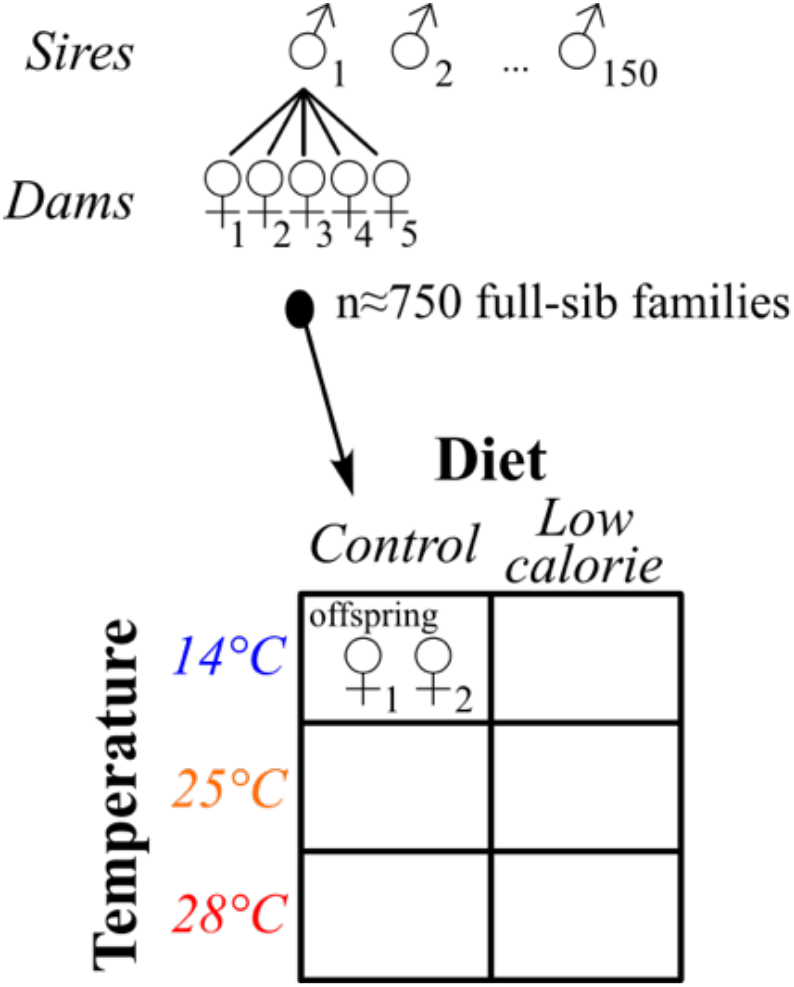
Experimental design. We mated 150 sires each to 5 dams, which we left to lay for 12 hours in each of 6 treatments. This replicated c.750 families in each of the 6 treatments. Two female offspring that developed in each treatment were then assayed for productivity (number of offspring produced), our proxy for fitness.

### Population establishment

We used a previous collection of *D. melanogaster* sampled at eight locations from discarded grape skins at wineries covering a variety of thermal environments across Victoria (Australia) in March-April 2021 (Walter *et al*. 2026a). Briefly, we collected c.50 inseminated females from each location and generated isofemale lines in the lab by allowing females to lay for 4 days in 40mL vials containing 7mL of standard yeast-potato-dextrose medium (**Appendix S2**). Larvae were left to develop at 25 °C on a 12:12 h light:dark cycle. We then generated eight mass-bred populations by combining five male and five female virgins from each isofemale line (c.50 lines per population) into 300mL plastic bottles containing 62.5mL of standard food. We maintained the mass-bred populations in 5 bottles per population at 25°C.

After 12 generations, we combined the eight mass-bred populations into a single large mass-bred population representing a large pool of genetic variation present within Victoria, and which could be created by gene flow that is common at distances <500km (Kennington *et al*. 2003). We divided 500 individuals (250 females and 250 males) from each of the eight populations equally among 20 bottles with 62.5mL of standard food (n=200 flies per bottle; total N=4000 flies). We then maintained the large mass-bred population in 20 bottles at 25 °C for three generations to allow them to mix thoroughly while keeping the population size high.

### Breeding design and experimental setup

To create the breeding design, we crossed individuals from the third generation of the large mass-bred population. We collected virgin adults and mated 150 sires each to five females in a nested half-sibling quantitative genetics breeding design (Lynch & Walsh 1998), resulting in 750 full-sibling families. To produce the families, we left each sire in a vial to mate with five females for two days. We then transferred each female to individual 40mL vials containing 7mL of their standard diet. To replicate the families in each of six treatments, we allowed females to lay for 12 hours, and then transferred them to new vials for their next treatment. In this way, we transferred each female to lay eggs in six different vials containing their treatment diet (standard or 25% dilution), after which we added eclosion cards and randomised vials into their treatment temperatures (14, 25 and 28 °C) in three controlled-temperature cabinets with 12:12 light:dark cycles. Vials were density controlled by removing eggs to leave 40-60 eggs per vial. The total number of families varied slightly across treatments because females did not lay in every treatment, we lost 7 sires that failed to mate, and several females escaped during transfers. However, in each treatment we measured ≥135 sires each mated to >1 dam (**Table S1**), which provides strong replication for estimating additive genetic variance. In total, we assayed 6,906 females from 143 sires and 688 dams.

### Assaying fitness

As our proxy for fitness, we estimated productivity by quantifying the total number of offspring produced by a mated female that laid eggs across their most fecund period. This combines fecundity of the female with offspring viability to capture fitness as the total surviving offspring produced (Mackay 1985; Nguyen & Moehring 2015). To measure productivity, we first collected and counted the emergent flies from each development vial, and then randomly selected two females to assay productivity. Each female selected for the productivity assay was held individually in vials with a randomly selected male until sexually mature (started laying eggs). We then allowed the female to lay eggs during the most fecund period in two subsequent vials (Nguyen & Moehring 2015). Initially, we started with three vials, but found this too logistically challenging given that we were assaying 1500 females per treatment. The length of time spent in each vial differed between diet and temperature treatments (**Table 1**), and was determined by balancing density (preventing high densities that could influence larval survival) while ensuring that we capture as much of the most fecund period for each female in each treatment. Pilot experiments suggested that females did not mature when held at their 14 °C temperature, and so we kept these females at 16 °C to ensure maturity while maintaining a cold treatment. To ensure that females were similar ages when productivity was assayed, we set up females at the same time following eclosion. This was more important in the low-calorie diet where there was larger variation in development time, which meant 2-3 sets of females were set up (**Table 1**).

**Table 1.** For each treatment, the time in days it took to develop, and then the time spent at each stage of the productivity assay. Maturing time is the time for females to start laying eggs, and then laying time is the time females spent laying in the productivity assay (in days). For treatments with large variation in development time, we set up several sets (last column) so that all females within a treatment were assayed at the same age.

| Treatment | Development time<br>(egg – eclosion) | Maturing time | Laying time | Number of sets |
| --- | --- | --- | --- | --- |
| 14°C, Standard diet | 35-46 | 8 | 11 | 2 |
| 14°C, Low calorie | 39-68 | 9 | 15 | 3 |
| 25°C, Standard diet | 8-12 | 5 | 6 | 1 |
| 25°C, Low calorie | 10-19 | 5 | 8 | 2 |
| 28°C, Standard diet | 7-11 | 5 | 6 | 1 |
| 28°C, Low calorie | 8-14 | 5 | 9 | 2 |

### Statistical methods

To quantify adaptive potential, we estimated the additive genetic variance in relative fitness, *V*_*A*_(*w*), across all treatments using a single multivariate generalised linear mixed model. We denote by *y*_*tijk*_ the number of offspring produced by the *k*th daughter assayed from the family of dam *j* nested within sire *i*, in treatment *t*. Counts were modelled as

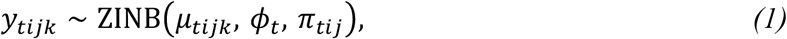

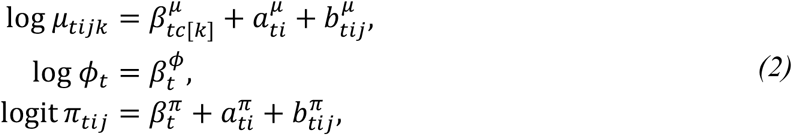

where the zero-inflated negative binomial (ZINB) is a two-component mixture with probability π_*tij*()_ that an observation is a structural zero, and otherwise it is drawn from a negative binomial distribution with mean µ_*tijk*_ and shape ϕ_*t*_ with variance µ + µ^1^/ϕ. The zero-inflation component captures zeros in excess of those expected under the negative binomial (c.26% of counts were zero, mostly on the low-calorie diet and at the cold temperature). These excess zeros represent whether a female reproduces at all (hereafter, reproductive failure), which may be distinct from the processes determining how many offspring each female produces. The intercepts of the mean were specific to each assay set *c* within treatment (11 combinations), while those of the zero-inflation term and shape were specific to treatment. We used the same random effects structure for each of µ and π as these two components jointly determine the expected fitness and its genetic variance. The components had random effects for sire 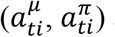 and dam 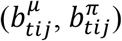 distributed as

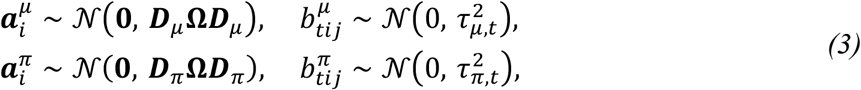

where ***D***_•_ = diag(***σ***_•,1_, …, ***σ***_•,6_) is the diagonal matrix of treatment-specific sire standard deviations, *τ*_•,*t*_ are the dam standard deviations, and *Ω* the shared 6 × 6 unstructured correlation matrix of sire effects across treatments.

The fixed effects for the mean and the shape were given flat priors, and the zero-inflation intercepts were given π_*t*_ *∼* Logistic(0,1), which is uniform on the probability scale. All standard deviations were given half-Student-*t* priors (df = 3, scale = 3.4), and the sire correlation matrix a uniform prior over correlation matrices, *Ω ∼* LKJ(1) (Lewandowski *et al*. 2009). The model was implemented directly in Stan (Carpenter *et al*. 2017) and fitted in R (R Core Team 2025) using *cmdstanr* (Gabry *et al*. 2024), running four chains of 6000 iterations including 2000 warmup, target acceptance probability 0.95 and maximum tree depth 12. We assessed convergence using the rank-normalised *R*[statistic (Vehtari *et al*. 2021), requiring 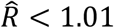 for all parameters, with bulk effective sample sizes above 400 and no divergent transitions. Model adequacy was assessed using posterior predictive checks.

In the ZINB model (equation 1), the expected fitness of a sire’s daughters is the product of the probability of not being an excess zero and the negative binomial mean, each averaged over assay sets and dam effects, so that the additive genetic variance in relative fitness *V*_*A*,’_(*w*) depends on the distribution of sire effects in both the µ and π components. Using Stein’s lemma (Stein 1981; de Villemereuil *et al*. 2016), the among-sire variance in expected relative fitness can be written as the weighted sum of 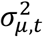 and 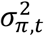 where the former has unit weight due to the log link and multiplicative decomposition (Bonnet *et al*. 2019). The latter is weighted by 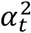, where *α*_*t*_, the average effect of 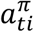 on relative fitness, is calculated numerically (see **Appendix S1**). Since for a nested half-sibling design, the sire variance is one quarter of the additive genetic variance (Lynch & Walsh 1998), the genetic variance in relative fitness *V*_*A,t*_(*w*), which represents adaptive potential, can be expressed as

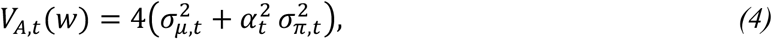

where –*q*_*t*_ ≤ *α*_*t*_ < 0 and *q*_*t*_ is the expected proportion of excess zeros in treatment *t*.

#### Interactive effects on the expression of additive genetic variance in fitness

We quantified interactions between temperature and diet by comparing the effect of diet on *V*_*A*_(*w*) across temperatures. Without an interaction, the effect of diet on genetic variance should be constant across temperatures. We therefore calculated the diet effect (difference in *V*_*A*_(*w*) between low-calorie and standard diets) at each temperature and compared it between each of the thermal extremes (14 °C and 28 °C) and the control temperature (25 °C). For the cold margin, the interaction is:

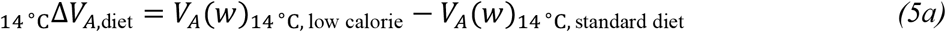

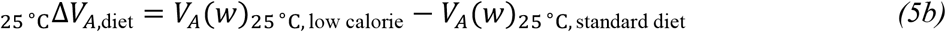

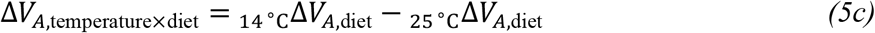

We evaluated equations 5a–5c at each posterior draw from the fitted model to obtain a posterior distribution of each contrast. To test our hypothesis that thermal stress strengthens the effect of low-calorie diet on genetic variance in relative fitness, we computed the posterior probability that the interaction effect is greater than zero. For the hot margin, we replaced 14 °C with 28 °C.

#### Interactive effects on the strength of additive genetic correlations in fitness

We tested whether temperature and diet interacted to influence additive genetic correlations in fitness, *r*_*A*_ (see **Appendix S1**). For each temperature margin, there are four treatments (two temperatures crossed with two diets) from which we calculated two contrasts. The first quantifies whether diet changes the genetic correlation across temperature,

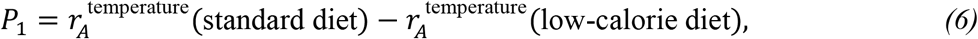

where, for the cold margin, 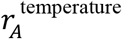 (standard diet) is the correlation between 14 °C and 25 °C on the standard diet. The second quantifies whether temperature changes the genetic correlation across diet,

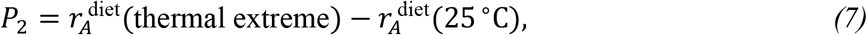

where 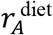 is the correlation between the standard and low-calorie diets at a given temperature (14 °C for the cold margin). If there is no interaction, both contrasts are expected to be zero. Positive values of *P*_4_ indicate that genetic correlations across temperature were weaker on the low-calorie diet, and positive values of *P*_1_ that genetic correlations across diet were stronger at the thermal extreme than at 25 °C. For each contrast we report the posterior mean, 90% HPD interval and posterior probability that it exceeds zero.

## Results

Mean fitness was highest on the standard diet at 25 °C and 28 °C and declined sharply on the low-calorie diet at every temperature, with the lowest fitness at 14 °C on both diets (**Fig. 2a**). Adaptive potential, quantified as *V*_*A*_(*w*), was lowest on the standard diet at 25 °C (0.186; 0.125–0.244 90% HPD) and 28 °C (0.126; 0.086–0.167) and increased under cold stress and under nutritional stress, but not under heat stress alone (**Fig. 2b; Table 2**). Posterior mean *V*_*A*_(*w*) was higher on the low-calorie diet at every temperature, by 1.8-fold at 25 °C, 2.3-fold at 14 °C and 4.4-fold at 28 °C (**Fig. 3a**). The diet effect was largest at the cold margin, where the low-calorie diet increased *V*_*A*_(*w*) by 0.892 (0.090–1.597; P(>0) = 0.987), compared with 0.145 at 25 °C (−0.067–0.364; P(>0) = 0.871) and 0.434 at 28 °C (0.076–0.777; P(>0) = 0.995; **Fig. 3a**). Genetic variation in reproductive failure contributed most at 14 °C where failure was common, accounting for c.25% and c.31% of *V*_*A*_(*w*) at 14 °C on the standard and low-calorie diets, compared with 10% or less in the other treatments (**Fig. 2c**). Relative to the control temperature, the diet effect at 14 °C was larger by 0.747 (−0.040–1.546), with a posterior probability of 0.955 that this interaction was positive (equations 5a–c; **Fig. 3a**). This is consistent with our hypothesis that cold stress strengthens the effect of nutritional stress on genetic variance in fitness. At the hot margin, the interaction was smaller and less certain (0.289; -0.133–0.704; P(>0) = 0.880). Although hot and cold thermal stress both amplified the expression of genetic variance in response to nutritional limitation, cold stress had a stronger interactive effect with diet stress.

**Table 2.** Matrix showing additive genetic variance in relative fitness, *V*_*A*_(*w*) (diagonal, grey shading) and additive genetic correlations in fitness across treatments (off-diagonal). 90% HPD intervals are in brackets. Above the diagonal is the posterior probability that the larger of the two genetic variances (the higher value on the diagonal) exceeds the smaller.

|  | 14°C,<br>Standard | 14°C,<br>Low<br>calorie | 25°C,<br>Standard | 25°C,<br>Low<br>calorie | 28°C,<br>Standard | 28°C,<br>Low<br>calorie |
| --- | --- | --- | --- | --- | --- | --- |
| 14°C,<br>Standard | 0.710<br>(0.405,<br>0.985) | 0.987 | >0.999 | 0.961 | >0.999 | 0.707 |
| 14°C,<br>Low<br>calorie | 0.799<br>(0.659,<br>0.949) | 1.602<br>(0.873,<br>2.316) | >0.999 | >0.999 | >0.999 | 0.990 |
| 25°C,<br>Standard | 0.595<br>(0.409,<br>0.780) | 0.614<br>(0.419,<br>0.799) | 0.186<br>(0.125,<br>0.244) | 0.871 | 0.945 | 0.976 |
| 25°C,<br>Low<br>calorie | 0.466<br>(0.182,<br>0.728) | 0.421<br>(0.163,<br>0.703) | 0.434<br>(0.175,<br>0.708) | 0.331<br>(0.119,<br>0.529) | 0.965 | 0.823 |
| 28°C,<br>Standard | 0.541<br>(0.351,<br>0.737) | 0.497<br>(0.295,<br>0.688) | 0.847<br>(0.749,<br>0.954) | 0.465<br>(0.202,<br>0.739) | 0.126<br>(0.086,<br>0.167) | 0.995 |
| 28°C,<br>Low<br>calorie | 0.391<br>(0.114,<br>0.682) | 0.306<br>(0.004,<br>0.603) | 0.428<br>(0.159,<br>0.694) | 0.471<br>(0.151,<br>0.795) | 0.625<br>(0.410,<br>0.842) | 0.560<br>(0.218,<br>0.918) |

**Fig. 2.**
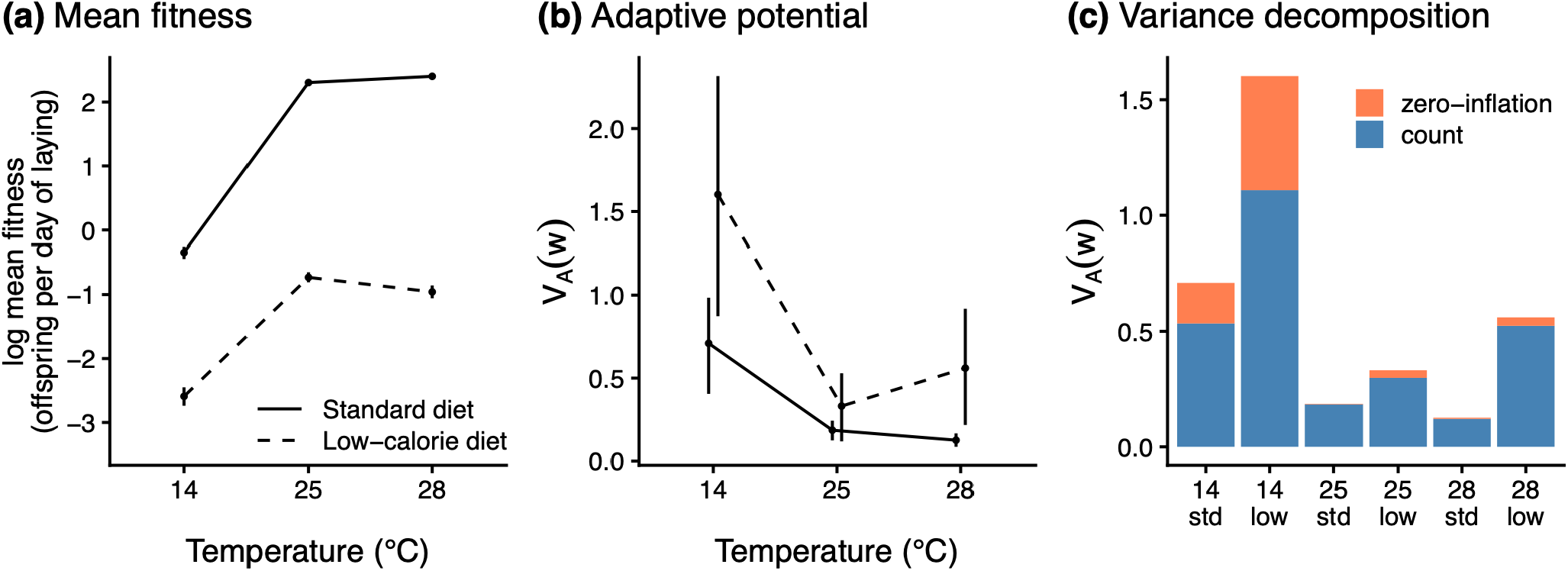
Mean fitness and adaptive potential across combinations of temperature and diet. **(a)** Mean fitness in each treatment, as the log of the expected number of offspring produced per female per day of laying. **(b)** Additive genetic variance in relative fitness, *V*_*A*_(*w*), the measure of adaptive potential. In **(a)** and **(b)**, points are posterior means and bars 90% highest posterior density (HPD) credible intervals, most of which are smaller than the points in **(a)**. Solid lines represent the standard diet, and dashed lines the low-calorie diet. **(c)** Posterior mean *V*_*A*_(*w*) in each treatment (std = standard diet; low = low-calorie diet), divided into the contributions of sire variation in the number of offspring produced by reproducing females (count component; blue) and in reproductive failure (zero-inflation component; orange), following equation 4.

**Fig. 3.**
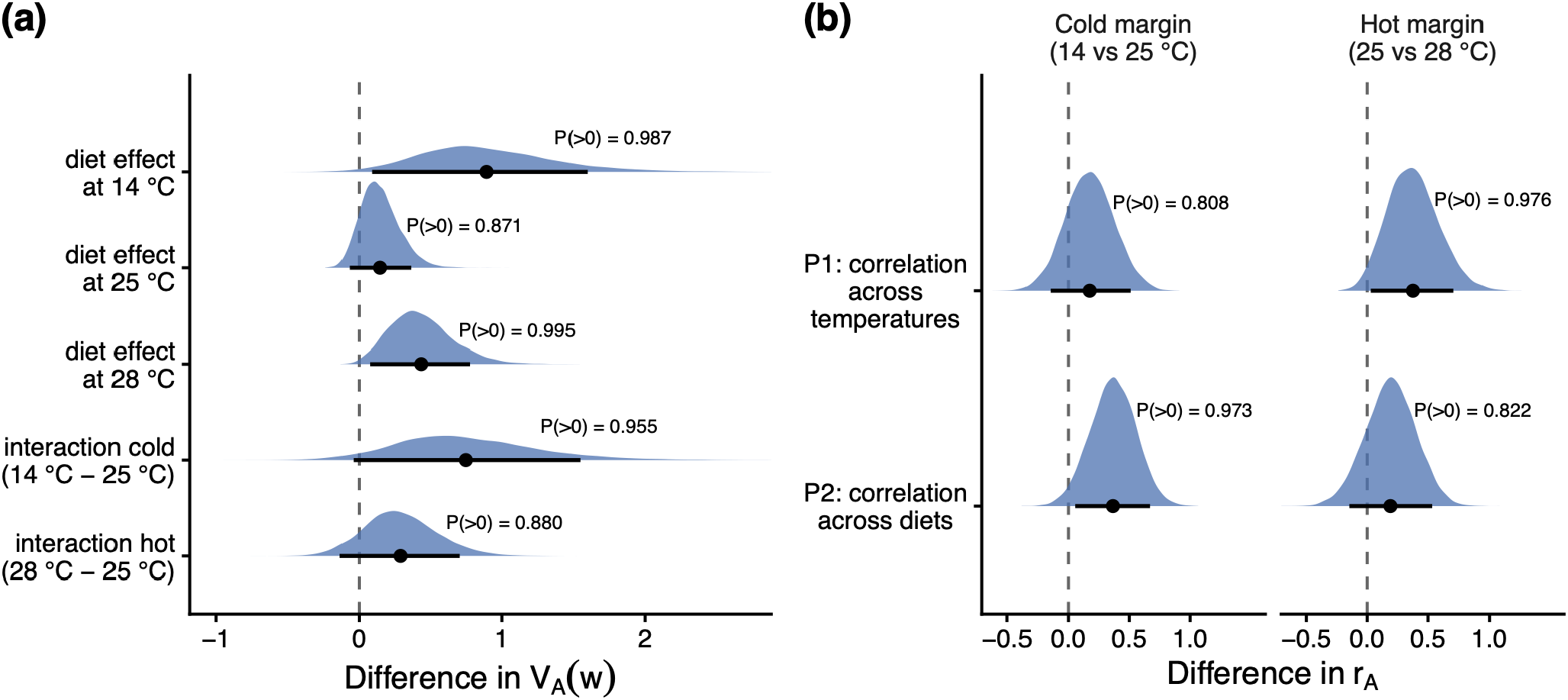
Interactions between temperature and diet on genetic variance and genetic correlations in fitness. **(a)** Posterior distributions of the effect of the low-calorie diet on *V*_*A*_(*w*) at each temperature (low-calorie minus standard diet), and of the temperature × diet interactions at the cold and hot margins (the diet effect at the thermal extreme minus that at 25 °C; equations 5a–c). The thin upper tails of the diet effect at 14 °C and of the cold-margin interaction extend beyond the plotted range. **(b)** Posterior distributions of the two correlation contrasts at the cold (14 vs 25 °C) and hot (25 vs 28 °C) margins (equations 6–7). *P*_1_ is the genetic correlation across temperatures on the standard diet minus that on the low-calorie diet; positive values mean that nutritional stress weakened the correlation across temperatures. *P*_2_ is the genetic correlation across diets at the thermal extreme minus that at 25 °C; positive values mean that genotypes ranked more consistently across diets at the thermal extreme. In both panels, points are posterior means and bars 90% HPD intervals; the dashed line marks zero, the value expected if temperature and diet act independently (and, for the diet effects in (**a**), if diet has no effect).

Genetic correlations in fitness between treatments were all positive (*r*_*A*_ = 0.306–0.847), suggesting that for the most part, sire performance in one treatment predicted sire performance in other treatments (**Table 2**). On the standard diet, genetic correlations across temperatures were strong (**Table 2**: *r*_*A*_ 25 °C, 28 °C = 0.847; 14 °C, 25 °C = 0.595; 14 °C, 28 °C = 0.541), which contrasted with weaker correlations across temperatures on the low-calorie diet (**Table 2**: *r*_*A*_ 25 °C, 28 °C = 0.471; 14 °C, 25 °C = 0.421; 14 °C, 28 °C = 0.306). Genotype performance was therefore less consistent across temperatures when nutrition was limited.

Comparing the strength of correlations across the four treatments at each thermal margin provided evidence that temperature and diet interacted to determine genetic correlations in fitness (equations 6–7; **Fig. 3b**). At the hot margin, the genetic correlation between 25 °C and 28 °C was weaker on the low-calorie diet than on the standard diet (*r*_*A*_ = 0.471 versus 0.847; *P*_4_ = 0.376; 0.029–0.709 90% HPD; P(>0) = 0.976), whereas the corresponding change at the cold margin was in the same direction but weakly supported (*r*_*A*_ = 0.421 versus 0.595; *P*_4_ = 0.174; -0.146–0.511; P(>0) = 0.808). Conversely, the genetic correlation between diets was stronger at 14 °C than at 25 °C (*r*_*A*_ = 0.799 versus 0.434; *P*_1_ = 0.366; 0.055–0.670; P(>0) = 0.973), and was in the same direction but weakly supported at 28 °C (*r*_*A*_ = 0.625 versus 0.434; *P*_1_ = 0.191; -0.147–0.534; P(>0) = 0.822). Temperature and diet therefore interacted to determine how consistently sires ranked across environments, with different effects at each margin: at the hot margin, nutritional stress made a sire’s performance at one temperature a poorer predictor of its performance at the other, whereas at the cold margin, the cold treatment made a sire’s performance on one diet a better predictor of its performance on the other. **Fig. 4** illustrates how weakening genetic correlations influence genotype rankings across temperature and diet. Sires largely kept their rank across a change in temperature alone (*r*_*A*_ = 0.847), but changed rankings more frequently across temperature combined with diet (*r*_*A*_ = 0.428) and across the temperature extremes under nutritional stress (*r*_*A*_ = 0.306; **Fig. 4**).

**Fig. 4.**
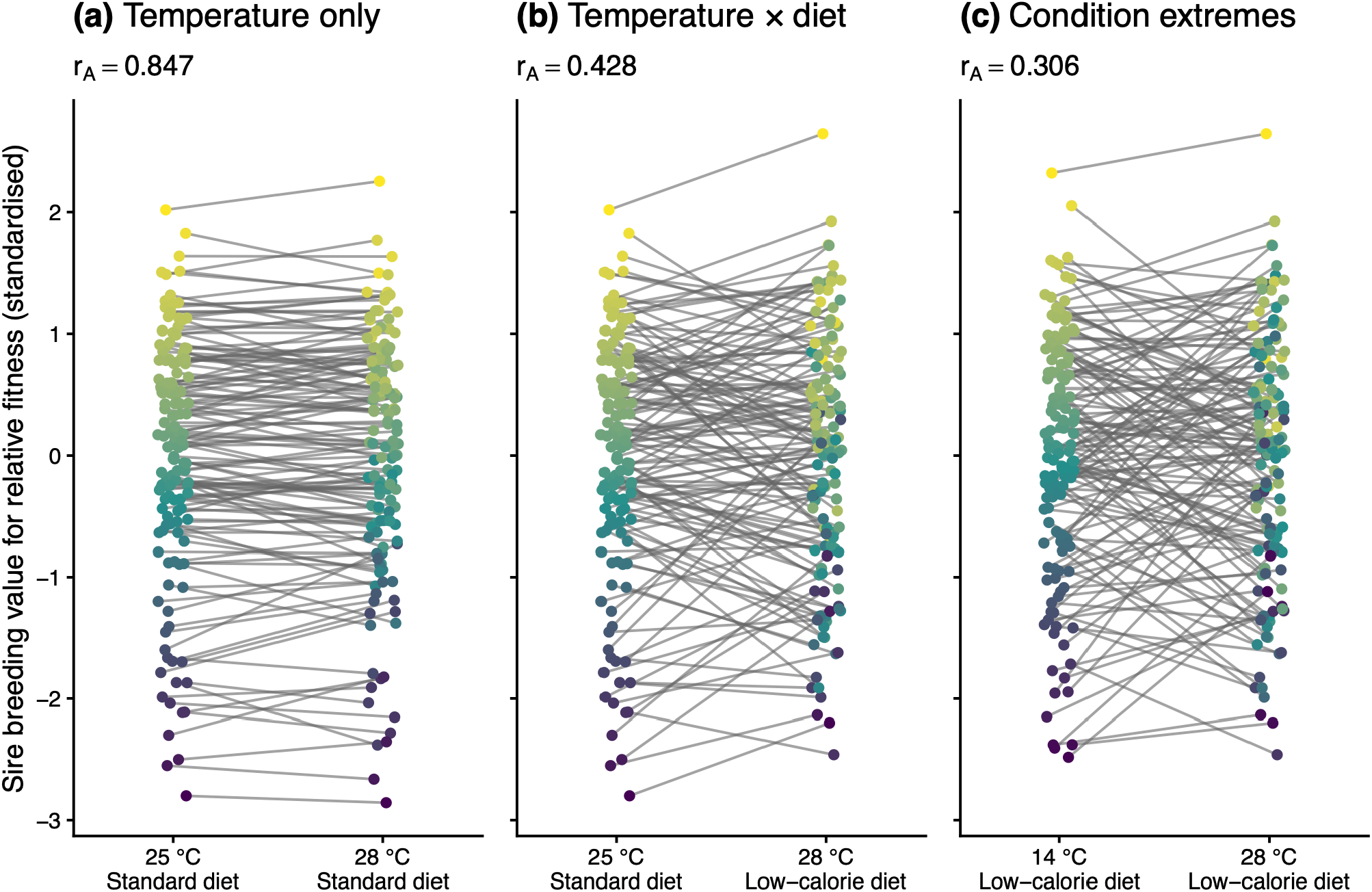
Illustration of the biological meaning of strong and weak genetic correlations across environments. Reaction norms show posterior mean sire breeding values for relative fitness, standardised within each environment, for three pairs of environments. Colours are ordered by sire breeding value in the left-hand environment, so weaker correlations appear as more re-shuffling of colours (sire rank) across environments. **(a)** Environments differing only in temperature (25 °C versus 28 °C, standard diet): sire rankings are largely conserved, reflecting a strong genetic correlation (*r*_*A*_ = 0.847). **(b)** Environments differing in both temperature and diet (25 °C standard diet versus 28 °C low-calorie diet): sires change rank more often (*r*_*A*_ = 0.428). **(c)** The temperature extremes on the low-calorie diet (14 °C versus 28 °C): the weakest correlation among all pairs of treatments (*r*_*A*_ = 0.306). The value above each panel is the additive genetic correlation in fitness (*r*_*A*_; **Table 2**).

## Discussion

Predicting whether populations can adapt rapidly enough to avoid extinction under global change requires understanding how adaptive potential itself changes across complex environments. Using a nested half-sibling breeding design with a large, outbred population of *Drosophila melanogaster*, we show that additive genetic variance in fitness is strongly environment-dependent and shaped by interactions between temperature and diet. Genetic variance in fitness increased under nutritional stress, most markedly at the cold margin (**Figs 2b, 3a**; **Table 2**). Temperature and diet also interacted to determine genetic correlations between treatments: at the hot margin, the correlation between temperatures weakened under nutritional stress, whereas at the cold margin, the correlation across diets strengthened under cold conditions compared to 25 °C (**Fig. 3b; Table 2**). These results suggest that adaptive potential under combined environmental stressors may not be reliably predicted from single stressors in isolation, and that the interactive effects of stressors influence adaptive potential differently at hot versus cold margins. Understanding the capacity for populations to adapt to global change therefore needs to consider how cryptic genetic variation and genotype-by-environment interactions determine adaptive potential across multidimensional environmental change.

### Release of cryptic adaptive potential under stress

Our findings are consistent with theoretical predictions that environmental perturbation can release cryptic genetic variation that is hidden under benign conditions (Hermisson & Wagner 2004; Paaby & Rockman 2014). Stabilising selection is expected to erode additive genetic variance under benign conditions, fixing phenotypes around locally optimal trait combinations. When organisms experience novel or stressful environments, previously neutral, conditionally expressed, or epistatic variation could be converted into additive genetic variance, increasing adaptive potential (Blows & Sokolowski 1995; Hoffmann & Merilä 1999). Genetic variance in fitness was higher on the low-calorie diet at every temperature, most clearly at 14 °C and 28 °C, with the largest increase at the cold margin. This pattern suggests that combined cold and nutritional stress disrupts developmental or physiological buffering mechanisms, exposing genetic differences among genotypes that are not expressed under benign conditions. Increased adaptive potential at both margins (particularly the cold margin) suggests that cryptic genetic variation expressed under combined stress cannot be inferred from single-stressor treatments. Cryptic adaptive potential may therefore be underestimated in standard laboratory assays that manipulate environmental axes independently.

The contrasting responses at the two thermal margins, with the largest increase in genetic variance under cold and nutritional stress but greater re-ranking of genotypes across temperatures under heat and nutritional stress, may reflect differences in the physiological basis of thermal tolerance. Heat tolerance often depends on relatively conserved pathways, including heat shock protein expression, membrane stabilisation and protein homeostasis (Milton *et al*. 2003), which are likely subject to strong stabilising selection that removes additional segregating variation that would otherwise be expressed under heat stress. Lower cryptic variation within a limited set of pathways could also explain why, under heat and nutritional stress, the increase in genetic variance relative to the control temperature was smaller than at the cold margin, whereas genotypes showed more re-shuffling in their relative performance (i.e., weaker genetic correlations across temperatures). Cold tolerance, by contrast, can draw on a broader range of mechanisms, such as metabolic depression, membrane lipid remodelling, developmental timing and energy allocation (Overgaard & MacMillan 2017), which is consistent with the idea that traits with a broader genetic and physiological basis show greater environmental sensitivity and scope for releasing cryptic genetic variance (Houle 1992; Sgrò & Hoffmann 1998). Slowed development under combined cold and nutritional stress may further prolong larval exposure to stress, amplifying the expression of genetic variance (King *et al*. 2022). These mechanistic interpretations require direct testing, but are consistent with the idea that different combinations of environmental extremes vary in their capacity to release cryptic genetic variation. Regardless of mechanism, populations at cold geographical margins, which may already face nutritional stress due to shorter growing seasons or reduced prey availability, could harbour substantially greater adaptive potential than would be detected by single-stressor assays. This is consistent with evidence that cold-edge populations show elevated adaptive capacity and stronger patterns of local adaptation (Hargreaves & Eckert 2019).

### Higher-order G×E and the multidimensional nature of adaptive potential

Our results represent higher-order genotype-by-environment interactions (G×E×E), in which both the amount of genetic variance and the extent of genotype re-shuffling across environments depend on interactions between environmental axes rather than any single axis alone. Such higher-order interactions are likely to be common in nature where organisms simultaneously experience variation in temperature, resource availability, species interactions, and other abiotic and biotic factors (Des Marais *et al*. 2017; Hudak & Dybdahl 2023). Ecological research has often demonstrated that multiple stressors interact synergistically or antagonistically at the phenotypic and demographic levels (Folt *et al*. 1999; Crain *et al*. 2008; Piggott *et al*. 2015). Experimental manipulation has also shown that temperature and diet can influence the expression of genetic variance in body size and life history traits (de Moed *et al*. 1997; Choy *et al*. 2024). Our results extend this to fitness, showing that combinations of stressors can shape both the expression of additive genetic variance in fitness and which genotypes are favoured.

Classic models of evolutionary rescue assume constant additive genetic variance and that environmental change can be described along a single axis that shifts the location of the fitness optimum (Gomulkiewicz & Holt 1995; Bell & Gonzalez 2009; Chevin *et al*. 2010). Under these assumptions, adaptive potential is approximated from genetic variance measured in a single novel environment. Our findings challenge both assumptions. Additive genetic variance for fitness and genetic correlations across environments changed substantially in response to combined stress in ways that were not always predictable from single-stressor environments. If such higher-order G×E interactions are common, forecasts of evolutionary rescue based on simplified environmental manipulations are likely to underestimate or misrepresent adaptive capacity of populations in response to realistic, multidimensional environmental change.

### Genetic correlations and adaptive potential

Genetic correlations across environments reveal the extent to which the same genotypes are favoured in different environments (Falconer 1952; Via & Lande 1985). Negative genetic correlations, where different genotypes are favoured across environments, may be rare and correlations may often remain positive but vary in magnitude (Sgrò & Hoffmann 2004; Saltz *et al*. 2018). We found positive but highly variable genetic correlations across treatments (**Table 2**), and at the hot margin, nutritional stress weakened the genetic correlation across temperatures (**Fig. 3b**). Therefore, while the same genotypes are likely to be favoured across environments when populations face moderate thermal variation under benign nutritional conditions (*r*_*A*_ = 0.847), combinations of temperature and diet stress showing the weakest correlations (*r*_*A*_ = 0.306) suggest that the genotypes best suited to cold nutritional stress are not always the same as those best suited to heat with nutritional stress (**Fig. 4c**). For populations facing both increasing temperatures and reduced nutritional quality, a realistic scenario under global change, weak positive genetic correlations mean that adapting to different combinations of stress could require different genotypes, reshaping adaptive potential without negative genetic correlations or a large increase in genetic variance.

### Adaptive potential in response to combinations of stressors

Our focus on adaptive potential in response to temperature and diet only captures two of the many possible environmental variables in the wild. In reality, global change will involve many more interacting environmental factors (Folt *et al*. 1999; Crain *et al*. 2008; Todgham & Stillman 2013). In particular, it would be valuable to understand how changes in temperature and diet determine adaptive potential in the presence of changes in biotic factors, such as inter- and intra-species interactions, immune activation, or the presence of predators or parasites (Traill *et al*. 2010; Tseng & O’Connor 2015; Hector *et al*. 2020). It is also likely that specific diet shifts could interact with temperature to determine the expression of genetic variation (Chakraborty *et al*. 2020; Choy *et al*. 2024). To accurately predict the adaptive capacity of natural populations under global change, we therefore need to better understand how realistic combinations of stressors influence adaptive potential and population persistence across generations.

### Limitations for inferring adaptation

Although additive genetic variance in relative fitness provides a measure of adaptive potential, its ability to predict long-term evolutionary outcomes is likely to be limited. The expression of additive variance following environmental change may partly reflect the conversion of non-additive (dominance or epistasis) variance into additive variance, which may be transient when recombination in the following generations breaks up combinations of alleles underlying the increase in adaptive potential (Barton & Turelli 1987), resulting in a subsequent decline in adaptive potential. Furthermore, theoretical work highlights the restrictive assumptions required for short-term genetic variance to accurately predict long-term adaptation (Frank & Slatkin 1992; Ewens 2024). This is in part because adaptation depends not only on adaptive potential, but also on demographic parameters. Environments that maximise additive genetic variance may simultaneously depress mean fitness to levels that constrain effective population size and increase the probability of extinction. This means that if combined stressors reduce mean fitness below the threshold required for population persistence, demographic collapse may occur before adaptive evolution can restore positive growth (Lande 1988; Gomulkiewicz & Holt 1995; Bell 2017). Adaptive potential and demographic robustness must therefore be considered jointly to predict outcomes for the population. Experimental evolution under combinations of stressors would allow direct tests of whether environments that initially expose high additive variance are those in which adaptation proceeds most rapidly, and whether such adaptation is sufficient to prevent extinction.

## Conclusions

Our results show that adaptive potential is itself environmentally dependent and shaped by interactions among stressors. Additive genetic variance for fitness increased under nutritional stress, most of all at the cold margin. Temperature and diet also interacted to determine genetic correlations across environments, showing that stressor combinations can reshape adaptive potential by altering which genotypes are favoured. Consequently, adaptive potential under combined environmental change may not be reliably inferred from single-stressor experiments. If interactions among stressors commonly expose cryptic genetic variation, current forecasting frameworks are likely to underestimate both the context in which adaptive capacity emerges, as well as the adaptive potential of natural populations to global change.

## Acknowledgments

GMW is grateful to have been supported by Australian Research Council fellowships DE200101019 and FT240100466. CMS was supported by Australian Research Council Discovery Projects DP220102958 and DP240102830. LY was supported by the Australian Research Council Centre of Excellence for Plant Success in Nature and Agriculture (CE200100015).

## Supplementary material for

**Table S1.** Replication of sires and dams in the productivity assay. Grey shading denotes the sires mated to >1 dam, with subtotals provided for only these cells.

| Treatment | Number of dams measured per sire |  |  |  |  |  | Total sires with >1 dam |
| --- | --- | --- | --- | --- | --- | --- | --- |
|  | 0 | 1 | 2 | 3 | 4 | 5 |  |
| 14°C, Standard diet | 12 | 1 | 7 | 21 | 58 | 51 | 137 |
| 14°C, Low calorie | 12 | 0 | 8 | 13 | 58 | 59 | 138 |
| 25°C, Standard diet | 12 | 1 | 3 | 15 | 48 | 71 | 137 |
| 25°C, Low calorie | 12 | 2 | 4 | 20 | 44 | 68 | 136 |
| 28°C, Standard diet | 7 | 1 | 1 | 7 | 39 | 95 | 142 |
| 28°C, Low calorie | 12 | 3 | 9 | 23 | 50 | 53 | 135 |

## Appendix S1 Additive genetic variance in relative fitness

In the ZINB model, expected fitness depends on both the count and zero-inflation components and thus the additive genetic variance in fitness comprises contributions from both. For a single sire, we stack the random effects of the two components across the six treatments, ***a*** = (***a***^µ ⊤^, ***a***^π⊤^)^⊤^, where 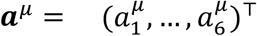 and likewise for ***a***^π^. From equation 3 of the main text, ***a*** *∼ N*(***0, S***) with

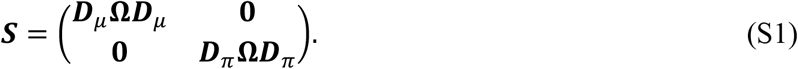

In treatment *t*, the expected fitness of a sire’s daughters, averaged over the dam effects 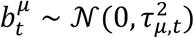 and 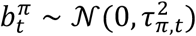, is

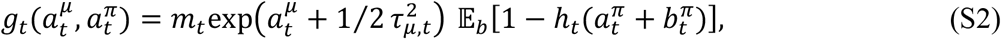

where *h*_*t*_(*x*) = logistic(*ψ*_*t*_ + *x*) is the probability of a structural zero, and 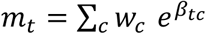 averages the set effects with weights *w*_+_ equal to the proportion of samples in each set. Mean fitness is 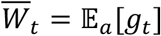 and 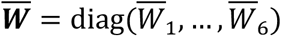.

The additive genetic variance in fitness is the variance of its least-squares regression on the sire effects (de Villemereuil *et al*. 2016). Given the sire effects are normally distributed, the coefficients of this regression are, by Stein’s lemma (Stein 1981), the average of the fitness gradient, E[∂*g*_*t*_/ ∂***a***] (Lande & Arnold 1983; Hansen & Houle 2008). Collecting these in the 12 × 6 matrix ***Ψ***, with Ψ_*kt*_ = E[∂*g*_*t*_/ ∂*a*_*k*_], the additive covariance matrix of fitness among sires is ***Ψ***^⊤^***S Ψ***. We multiply this by four for the half-sib design and divide by the mean fitness to give the additive genetic covariance matrix of relative fitness,

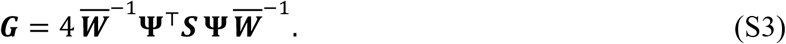

Since *g*_*t*_ depends only on 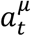 and 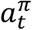, ***Ψ*** is a stack of two diagonal 6 × 6 blocks, one per component. For the count component, 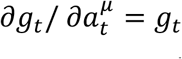, so the top block is 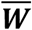 (Bonnet *et al*. 2019). For the zero-inflation component, 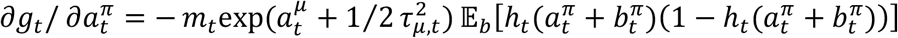. Since 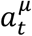 and 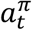 are independent and normal, the expected gradient, divided by *W*_’_, can be expressed as

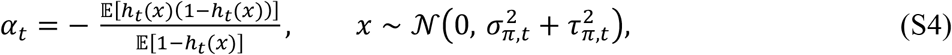

so that the bottom block is 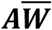, with ***A*** = diag(*α*_1_, …, *α*_6_), *α*_*t*_ < 0 for all *t*. We evaluated (S4) at every posterior draw using numerical quadrature.

Substituting 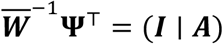 and (S1) into (S3) gives

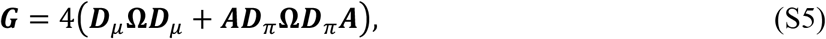

whose diagonal is equation 4 of the main text. Because 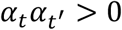, both components contribute to each covariance with the sign of 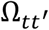. The corresponding genetic correlations are 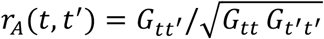.

## Appendix S2 Standard diet

The standard yeast-potato-dextrose medium contained potato flakes 18.2g/L; dextrose 27.3g/L; Brewer’s yeast 36.4g/L; agar 6.4g/L, Nipagin [10% w/v in ethanol] 10.9mL/L; and propionic acid 4.6mL/L.

## Notes

https://doi.org/10.5281/zenodo.23463230

### Competing Interest Statement

The authors have declared no competing interest.

